# Pre- and post-flowering impacts of natural heat events on yield components in wheat

**DOI:** 10.1101/2023.05.08.539933

**Authors:** Najeeb Ullah, Brian Collins, Jack Christopher, Troy Frederiks, Karine Chenu

## Abstract

1. CONTEXT Wheat crops are highly sensitive to elevated temperatures and experience significant yield losses when short periods of heat occur at sensitive developmental phases.
2. OBJECTIVE This research aimed at quantifying wheat responses of grain yield and yield components to heat indicators in fluctuating field conditions.
3. METHODS The impacts of high temperature on yield and its components were assessed for 20-35 wheat lines in irrigated multi-environment trials over three years. Genotypes were cultivated using a novel photoperiod extension method adjacent to some conventional yield plots with different sowing dates. In the photoperiod-extension method, either single stems or plant quadrates were tagged at specific growth stages and hand-harvested at maturity, while conventional plots were mechanically harvested at maturity. The impact of heat was estimated for events occurring at different developmental stages and different temperature thresholds (26 to 35^○^C).
4. RESULTS The closest correlation between heat events and grain number was observed for a threshold temperature of 28^○^C pre-flowering. For individual grain weight, the correlation was statiscally the closest for threshold temperatures above 32°C post-flowering. In the tested environments, grain number was most sensitive to heat between 300 and 200^○^Cd before flowering. For each hot hour (T > 28^○^C) during this period, wheat genotypes lost an extra 0.25 grains spike^-1^ or 90 grains m^-2^ on average. Post-flowering heat reduced individual grain weight by 0.26 mg (at spike harvest) and grain yield by 2.44 g m^-2^ (at plot harvest) for every hot hour (T > 32^○^C) between 0 and 500^○^Cd after flowering. Heat impacts were the clearest for plants with synchronised phenology. In addition, results suggest that heat impacts can be quantified more reliably using finer time units (i.e. hot hours rather than days).
5. CONCLUSIONS Natural heat events strongly impacted grain number and individual grain weight of wheat for temperatures above 28-30°C in well-watered conditions. Reduction in grain number and individual grain weight were strongly associated with accumulated hot hours that occurred during 200-300°Cd before and 0-500°Cd after flowering, respectively.
6. IMPLICATIONS The findings from this study will assist improvement for crop modelling in response to heat, development of relevant phenotyping methods and selection of cultivars with better adaptation to warmer environments.

## 1. Introduction

Wheat (*Triticum aestivum* L.) is a major cereal crop, contributing 20% of the daily calorific and protein needs to the global food supply (Shiferaw et al., 2013). Wheat is highly sensitive to heat stress. For instance, a 5.6% grain yield reduction has been estimated for each 1°C increase in the atmospheric mean temperatures (Lobell and Field, 2007). Similarly, controlled environment studies show a 3–5% reduction in wheat grain yields for each 1°C rise in mean temperature >15°C (Gibson and Paulsen, 1999). Wheat crops across many parts of the world already experience frequent high temperatures (e.g. > 34°C) during grain development (Asseng et al., 2011; Talukder et al., 2014) with significant grain yield losses (Hatfield et al., 2011). Modelling studies have reported a significant increase in the frequency of heat events over recent decades, particularly during the grain-filling period (Ababaei and Chenu, 2020), with further increases projected (Collins and Chenu, 2021; Field et al., 2014; Lobell et al., 2015). Therefore, quantifying the impact of natural heat on crop yields is critical for developing management practices to sustain food production under changing climates (Collins and Chenu, 2021; Flohr et al., 2018; Zheng et al., 2012).

Grain yield losses in wheat are strongly influenced by the duration, intensity, and timing of the heat stress events (e.g. Djanaguiraman et al., 2014; Chenu and Oudin, 2019). The reproductive and grain-filling phases of wheat are highly sensitive to heat stress. For example, a single hot day (30°C/20°C day/night temperature) during the pollen tetrad or meiosis stage significantly inhibits pollen viability and translates into reduced grain number (Saini and Aspinall, 1982). Hence, late-flowering tillers (secondary tillers) that experience heat during gametogenesis produce significantly fewer grains per spike than main tillers that were subjected to heat at flowering (Aiqing et al., 2018).

Post-flowering heat reduces individual grain weight (IGW; Girousse et al., 2021, 2018), particularly during the early phases of grain development (0-10 days post-anthesis in Egli, 1998),; 0-15 days after anthesis in Stone and Nicolas, 1995). Although sustained high temperatures (30–38°C) from flowering to maturity can significantly limit wheat grain yield formation, the magnitude of the effect across reported experiments varied from 20 to 50% (Wardlaw et al., 1989; Tewolde et al., 2006). In contrast, controlled environment studies indicated that a brief heat event during the sensitive phase could significantly reduce the IGW of wheat (Talukder et al., 2014). For example, a single hot day (40/21°C day/night) was found to induce a 14% reduction in IGW (Stone and Nicolas, 1998), while in another study, 14 hot days (32/22°C) during early grain-filling reduced IGW by 44% (Djanaguiraman et al., 2020). However, reports of such detailed information are limited to controlled environments in parts due to difficulties to observe impacts of brief heat events of plants grown in the field. In addition, results from controlled environments do not systematically translate in the field (e.g. Rebetzke et al., 2014) due for instance to heat effects on pots and roots. Furthermore, pot studies commonly focus on extreme stress, sometimes screening for survival, although the environments targeted by breeders are very different, with stress patterns varying in time and intensity (e.g. Chenu et al., 2013; Collins and Chenu, 2021, Ababaei and Chenu, 2020). Heat response, and in particular threshold temperatures, have not been studied in detail, in naturally fluctuating field conditions and during specific developmental stages.

A limitation on screening for heat tolerance in field conditions is that observed heat response can be confounded by genotypes of different maturity types being impacted by heat event(s) at different developmental stages. The literature discussed above suggests that the same heat events in one trial hence likely result in different impacts on yield and yield components for genotypes of different maturity types. To reduce this potentially confounding effect, a new method has been developed to bring genotypes of different maturity types to a similar developmental stage during the post-flowering period in field heat tolerance trials using photoperiod extension (Ullah et al., 2023).

This study aimed to (i) characterise the most heat-sensitive developmental stage(s) of wheat crops for grain yield components in natural field conditions, (ii) determine threshold temperature (intensity and duration) for these developmental stages and (iii) quantify the impact of heat on grain yield components. Fully irrigated field experiments were conducted at three locations over three consecutive years, each with two sowing times. The impact of high temperature on grain yield components of wheat was quantified at spike and sub-plot (i.e. quadrats) levels with matched flowering using the photoperiod extension method (PEM) developed by Ullah et al. (2023). These results were compared to those obtained from adjacent conventional yield plots with natural flowering, as has been more commonly used in previous studies (Thistlethwaite et al., 2020). The finding could assist in (i) improving modelling capability to simulate the performance of genotypes in a wide range of environments to better predict impact of heat stress in different scenarios, and (ii) develop phenotyping method for relevant high-throughput screening of heat tolerance. Overall, we anticipate that results will assist in evaluating genotypes for improved adaptation to current and projected future environments.

## 2. Materials and methods

### 2.1. Planting materials

Wheat (*Triticum aestivum* L.) genotypes used in the study (Table S1, Supplementary) include commercially Australian cultivated cultivars such as Suntop, Spitfire, Gregory, Janz, Hartog, EGA Wylie, Corack, Yitpi, Mace and Scout. A set of CIMMYT genotypes described as heat tolerant under Australian environments was obtained from the University of Sydney (Thistlethwaite et al., 2020). The remaining wheat genotypes were selected from a multi-reference parent nested association mapping (MR-NAM) population developed for screening for heat and drought tolerance in wheat (Christopher et al., 2015, 2021; Richard, 2017; Fletcher, 2020). In total, 35 wheat genotypes were tested in PEM and plot trials, with 32 genotypes at each trial, except for the PEM trial with quadrat harvest in 2020, when only 20 selected genotypes were used (Table S1, Supplementary). In conventional plot trials, 32 genotypes were used in each trial.

### 2.2. Field trials

Field trials were conducted over three consecutive years, from 2018 to 2020, at three locations across south-eastern Queensland, Australia, at The University of Queensland Research Farm, Gatton (27°34′50″S, 152°19′28″E), Queensland Department of Agriculture and Fisheries Hermitage Research Station, Warwick (28°12’40’’S, 152°06’06’’E) and at the Tosari Crop Research Farm, Tummaville (27°49’09”S 151°26’15”E). A total of 35 wheat genotypes were tested under different growing environments (Table S1) using conventional plots and the newly developed PEM (Ullah et al., 2021, 2023). Each year, the trials were established in a randomised complete block design with two times of sowing and four replicates per genotype. To ensure a range of growing temperatures across developmental stages, trials were sown at dates ranging from late May to early September (Table 1). Standard crop management practices were adopted for all trials, including weed, disease and pest control. Experimental details are presented in Ullah et al. (2023). Briefly, under PEM, the genotypes were planted either in either single rows (2018 and 2019) or small plots (1×5m; 2020) (Table 1). At one end of each row or plot, artificial supplemented lights were set up, extending the photoperiod to 20h. The light intensity gradient induced a gradient in the flowering date along the test rows such that the photoperiod extension effect was greatest and the flowering date earliest, nearer to the lights (Ullah et al., 2023). For all genotypes, individual spikes and/or quadrats (0.5 linear meter) of each row or plot were tagged at anthesis (Zadoks growth stage 65; Zadoks et al., 1974) on a single day by selecting sections of the row at different distances from the lights for genotypes of different maturity type. Spike data were collected from approximately 20 individual spiks of each genotype tagged at anthesis. The plants were later harvested at maturity. Grain samples were manually counted to estimate grain number per unit of area and IGW.

**Table 1.** Trial characteristics, including the environment*, site, sowing date identifier (Time of Sowing, tSow), the types of the trial conducted (i.e. photoperiod-extension method (PEM) with tagging and harvesting of either single spikes or quadrates) and the sowing date. Also presented are mean (*Tmean*) and max (*Tmax*) daily temperature, day-time vapour pressure deficit (VPD) well as the mean duration of the pre- and post-flowering periods (“Days to flow.” and “Post flow. duration”, respectively). The duration of days to flowering and post-flowering were calculated from sowing to flowering and flowering to maturity, respectively. Also presented are the cumulative heat (days/hour) above 30^○^C during 0-500^○^Cd after flowering, and the heat environment type (HET) for the first tagging of trials with the PEM.

| Environment* | Site | Time of sowing | Trial type | Sowing date | $T_{mean}$ (°C) | $T_{max}$ (°C) | Mean VPD (kPa) | Days to flow. (days) | Post-flow. duration (days) | Cum. hours > 30°C (0-500°Cd after flow.) | Cum. days > 30°C (0-500°Cd after flow.) | HET*** |
| --- | --- | --- | --- | --- | --- | --- | --- | --- | --- | --- | --- | --- |
| GAT18-s1-T1 | Gatton | tSow1 | Spike & quadrate PEM | 03/07/2018 | 17.2 | 25.3 | 0.8 | 73 | 42 | 3 | 2 | HET1 |
| GAT18-s1-T2 | Gatton | tSow1 | Spike PEM | 03/07/2018 | 17.7 | 25.5 | 0.9 | 81 | 38 | 3 | 2 | HET1 |
| GAT18-s1-T3 | Gatton | tSow1 | Spike PEM | 03/07/2018 | 18.0 | 25.6 | 0.9 | 86 | 37 | 3 | 2 | HET1 |
| GAT18-s2-T1 | Gatton | tSow2 | Spike & quadrate PEM, & plots | 31/08/2018 | 21.0 | 28.5 | 1.1 | 53 | 39 | 65 | 12 | HET2 |
| GAT18-s2-T2 | Gatton | tSow2 | Spike PEM | 31/08/2018 | 21.9 | 29.0 | 1.1 | 60 | 35 | 67 | 13 | HET2 |
| GAT19-s1-T1 | Gatton | tSow1 | Spike PEM & plots | 09/07/2019 | 17.0 | 25.5 | 1.2 | 71 | 39 | 31 | 5 | HET2 |
| GAT19-s1-T2 | Gatton | tSow1 | Spike PEM | 09/07/2019 | 17.3 | 26.0 | 1.2 | 78 | 36 | 46 | 9 | HET2 |
| GAT19-s2-T1 | Gatton | tSow2 | Spike PEM & plots | 03/09/2019 | 20.3 | 31.0 | 1.8 | 59 | 35 | 129 | 20 | HET3 |
| GAT20-s1-T1 | Gatton | tSow1 | Quadrate PEM & plots | 26/05/2020 | 16.5 | 24.4 | 0.9 | 79 | 47 | 0 | 0 | HET1 |
| GAT20-s2-T1 | Gatton | tSow2 | Quadrate PEM & plots | 04/08/2020 | 20.0 | 29.0 | 1.2 | 65 | 37 | 45 | 14 | HET2 |
| TOS19-s1-T1 | Tosari** | tSow1 | Spike & quadrate PEM, & plots | 16/07/2019 | 18.1 | 27.2 | 1.1 | 79 | 36 | 45 | 12 | HET2 |
| TOS19-s1-T2 | Tosari** | tSow1 | Spike PEM | 16/07/2019 | 18.4 | 27.7 | 1.1 | 84 | 35 | 35 | 11 | HET2 |
| TOS19-s2-T1 | Tosari** | tSow2 | Spike PEM & plots | 06/09/2019 | 23.2 | 32.7 | 1.7 | 62 | 34 | 162 | 22 | HET3 |
| WAR18-s1-T1 | Warwick | tSow1 | Spike & quadrate PEM | 16/07/2018 | 16.0 | 24.0 | 0.9 | 73 | 39 | 1 | 1 | HET1 |
| WAR18-s1-T2 | Warwick | tSow1 | Spike PEM | 16/07/2018 | 16.7 | 24.5 | 0.9 | 79 | 37 | 7 | 2 | HET1 |
| WAR18-s2-T1 | Warwick | tSow2 | Spike & quadrate PEM | 12/09/2018 | 20.3 | 27.8 | 1.2 | 52 | 37 | 47 | 10 | HET2 |
| WAR18-s2-T2 | Warwick | tSow2 | Spike PEM | 12/09/2018 | 20.4 | 28.2 | 1.2 | 56 | 36 | 41 | 11 | HET2 |
| WAR20-s1-T1 | Warwick | tSow1 | Quadrate PEM & plots | 08/06/2020 | 14.8 | 23.3 | 0.7 | 101 | 49 | 0 | 4 | HET1 |
| WAR20-s2-T1 | Warwick | tSow2 | Quadrate PEM & plots | 12/08/2020 | 18.6 | 26.6 | 1.1 | 66 | 46 | 10 | 4 | HET1 |
| WAR20-s2-T2 | Warwick | tSow2 | Quadrate PEM | 08/06/2020 | 18.8 | 27.0 | 1.1 | 72 | 42 | 9 | 3 | HET1 |

\* An identifier of the site denominates environments as Gatton (GAT), Tosari (TOS) or Warwick (WAR); the year of the trial, the time of sowing (s1, tSow1 or s2, tSow2), and the tagging event (T1, T2, T3) when applicable.
\*\*While all other trials were fully irrigated, TOS19 tSow1 and TOS19 tSow2 only had supplementary pre-flowering irrigation and experienced mild post-flowering water stress.
\*\*\* Heat environment type 1 (HET1) corresponds to environments with 0 days or hours of temperature above of 32°C between 0 and 500°Cd after flowering, while heat environment type 2 (HET2) corresponds to 1-9 days or 1-49 hours of temperature > 32°C during the same period (Fig. S3). Heat environment type 3 (HET3) had more than 10 days or 50 hours of temperature > 32°C between 0 and 500°Cd after flowering (Fig. S3).

Conventional field plots were established adjacent to the PEM trials under similar management. At each location, these plots were sown on the same day as the PEM trials with two sowing dates and four replication plots per genotype, except in 2018, when only sowing 2 (tSow2) plots were established at Gatton. The plot size was 2×6 m in 2018 and 2020 and 1×6 m in 2019. All conventional plots were planted with a 25 cm row spacing and a population density of 130 plants m^-2^. Conventional plots were harvested using a small plot machine harvester at maturity when grain moisture was approximately 11%. Grain samples were manually counted to estimate grain number and IGW.

The combinations of sites, years, flowering dates and tagging events correspond to 20 different “environments”, each providing a unique set of temperatures and timing of heat with respect to the developmental stages of each genotype. These environments were defined using an identifier of the site (Gatton, GAT; Tosari, TOS; and Warwick, WAR); the year of the trial, the time of sowing (tSow1 or tSow2) and when applicable, the tagging event (T1, T2, T3; Table 1). All trials were fully irrigated, except at Tosari in 2019 (TOS19) and grown under non-limiting fertiliser conditions. TOS19 crops had only pre-flowering supplementary irrigation and experienced mild post-flowering water stress. Trials were irrigated with a boom irrigator (conventional plots) and wobbler sprinklers (PEM plots). Standard crop management practices, including weed, disease and pest control were adopted during the season across all the trials.

### 2.3. Weather data

Wheater data were collected from the site weather stations (Campbell Scientific). Light sensors (Apogee SP-110 pyranometers) were installed at 1.5 m height to measure light interception. HMP60 (Vaisala INTERCAP®) probes were used to measure the air temperature (Tair) and relative humidity (RH) at 1.5 m above the ground. Thermal time was calculated in degree days (°Cd) using the following equation (Jamieson et al., 1995).

### 2.4. Statistical analysis

Data were analysed using the R programming language (R Core Team, 2018). Data were presented as the mean (all tested genotypes) of the replicated data for each genotype. Data were analysed separately for spikes tagged individually (at synchronised phenology for all genotypes using the PEM), quadrate harvests (sections of each row or plot of the PEM) and conventional plots (unsynchronised phenology of genotypes).

Heat indicators were defined as maximum daily temperature (*T*_max_), mean temperature (*T*_mean_) or number of days or hours above specific threshold temperatures (*T*_thresh_). Threshold temperatures of 26°C, 28°C, 30°C, 32°C and 35°C were tested. Linear correlations were examined between heat indicators and grain yield components for different crop developmental periods (from 600°Cd before flowering to 500°Cd after flowering) for each 100°Cd. Heat-sensitive phases were identified, where grain yield components most strongly responded to the temperature thresholds.

## 3. Results

### 3.1. A wide range of heat environments were tested

In total, 20 environments were studied, with plants harvested in different years, locations, from different sowing dates and tagging dates each experiencing different heat patterns during development (Table 1). Across these environments, grain yield components of the tested wheat genotypes responded strongly to heat stress during different developmental stages. For example, the reduction in grain number and IGW was associated with both pre-flowering (200-300^○^Cd to flowering) and post-flowering (0-500^○^Cd after flowering) temperatures. On average, crops from the later sowing (tSow2) experienced more heat than from the first sowing (tSow1), both in terms of average daily maximum (*T*_max_) and mean (*T*_mean_) temperatures (Fig. S1, Table 1). The warmer conditions experienced by tSow2 crops compared with their respective tSow1 crops accelerated the crop phenology, shortening both the vegetative and grain-filling periods (Table 1).

The number of hours or days with temperature above threshold levels varied across environments, particularly with reference to flowering time. On average, wheat genotypes experienced relatively little heat stress pre-flowering, with GAT19-s2 and TOS19-s2 and WAR20-s2-T1 being the only environments with days or hours of *T*_max_ > 32^○^C before flowering (Fig. S2). In contrast, a wide range of post-flowering (0-500^○^Cd) heat was observed, with 1-21 days or 1-116 hours above the threshold temperature of 32^○^C (Fig. S3, Table 1). For simple comparisons, we classified growing environments based on post-flowering heat into three heat environment types. Heat environment type 1 (HET1) corresponds to environments with 0 days or hours of temperature > 32^○^C between 0 and 500^○^Cd after flowering, while heat environment type 2 (HET2) corresponds to 1-9 days or 1-49 hours of temperature > 32^○^C during the same period (Fig. S3). Heat environment type 3 (HET3) had more than 10 days or 50 hours of temperature > 32^○^C between 0 and 500^○^Cd after flowering (Fig. S3).

### 3.2. The impact of pre-flowering heat on grain number differed with heat environments

Despite fewer pre-flowering (200-300^○^Cd before flowering) hot days or hours in the tested environments (Fig. S2), the grain numbers responded clearly to heat indicators, particularly at the spike level and, to a lesser extent, at the plot level (Fig. 1). Part of these responses would be due to the impact of increasing temperature on the shortening of the vegetative growth period (Table 1), and thus the reduced duration to accumulate biomass pre flowering. The number of grains produced by studied genotypes also varied widely across the tested environments ranging from 26 to 36 grains spike^-1^ and from 2,500 to 12,500 grains m^-2^ at the plot level (Fig. 1). In this study, grain number was more sensitive to heat stress during 200-300^○^Cd before flowering (Fig.1), compared with any of the other 100°Cd intervals of development from between 600°Cd before flowering to flowering (data not shown). Significant but weaker correlations were also found between grain number and the other periods between 400^○^Cd pre-flowering to flowering (data not shown).

**Fig. 1.**
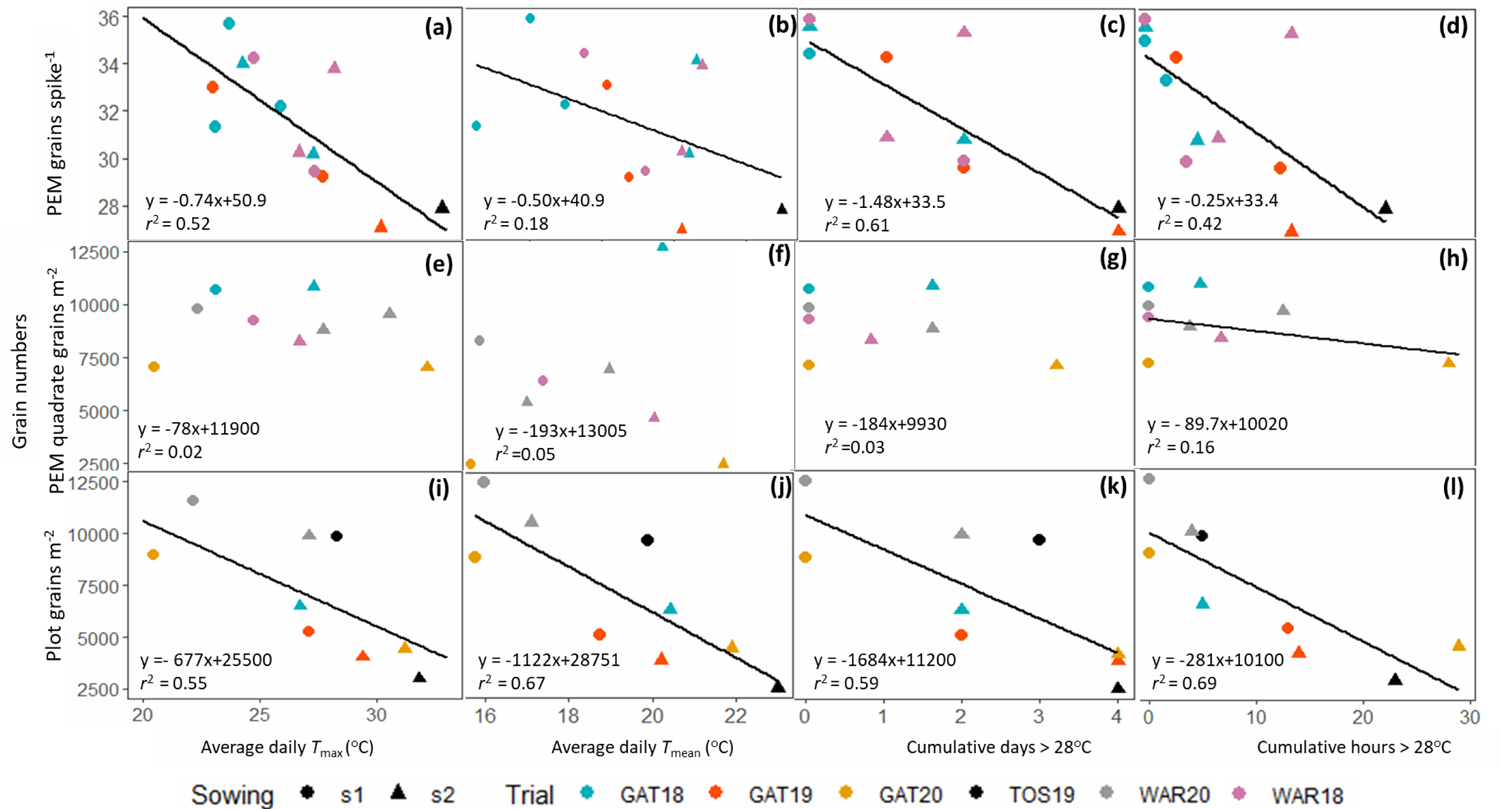
Changes in grain number in response to pre-flowering (a, e, i) daily maximum temperature (*T*max), (b, f, j) mean temperature (*T*mean)), cumulative (c, g, k) days and (d, h, l) hours above 28^○^C; all temperature indices were calculated between 300 and 200^○^Cd before flowering. Data correspond to the mean of 20 – 32 genotypes with four independent replicates in each environment. Spike data are collected from the individual spikes (∼20 for each replicate) exposed to heat at synchronised developmental stages. Quadrate data were collected by harvesting 0.5 m linear meter of plants tagged at synchronised flowering in the PEM. Plot data were collected from the whole conventional plots of naturally flowering genotypes (stage not synchronised during heat events). Quadrate and plot data are presented per unit area (m^-2^). Lines were plotted for regressions between mean grain number and temperature indicators that had a regression slope significantly different from zero (P<0.05).

For each 1^○^C increase in *T*_max_ between 300 to 200^○^Cd before flowering, genotypes produced 0.74 fewer grains spike^-1^ (PEM spike harvest, *r^2^* = 0.52, Fig. 1a) and 677 few grains m^-2^ at plot level (*r^2^* = 0.55, Fig. 1 i). With the PEM, grain number per spike responded relatively more strongly to *T*_max,_ and the number of hot hours than to *T*_mean_ (Fig. 1 a-d). For each hot day (*Tmax* > 28^○^C), between 300 and 200^○^Cd before flowering, wheat genotypes produced 4.4% and 15% fewer grains at the spike and plot levels, respectively (Fig. 1c, k). For each hot hour (> 28^○^C), grain number loss was estimated to be 0.75% and 2.8% in the PEM spikes and conventional plots, respectively, (Fig. 1 d, l). At the plot level, grain number response had similar coefficient of determination for the different heat indicators, ranging between 0.55 and 0.69 (Fig.1 i-l). Under quadrate harvest, the grain numbers response to heat indicators was relatively weak, possibly due to the absence of extremely hot (HET3) environments (Fig. e-h).

### 3.3. The impact of post-flowering heat on individual grain weight and yield also varied between heat environments

In the tested conditions, wheat plants produced the largest grains when there were no days with maximum temperature above 32^○^C i.e. for tSow1 of 2018 and 2020 (HET1, Fig. 2 and S3). Under such relatively low-stress temperatures, IGW of 44.1, 41.2 and 32.9 mg was observed for the PEM spikes, PEM quadrat, and plot harvests, respectively.

**Fig. 2.**
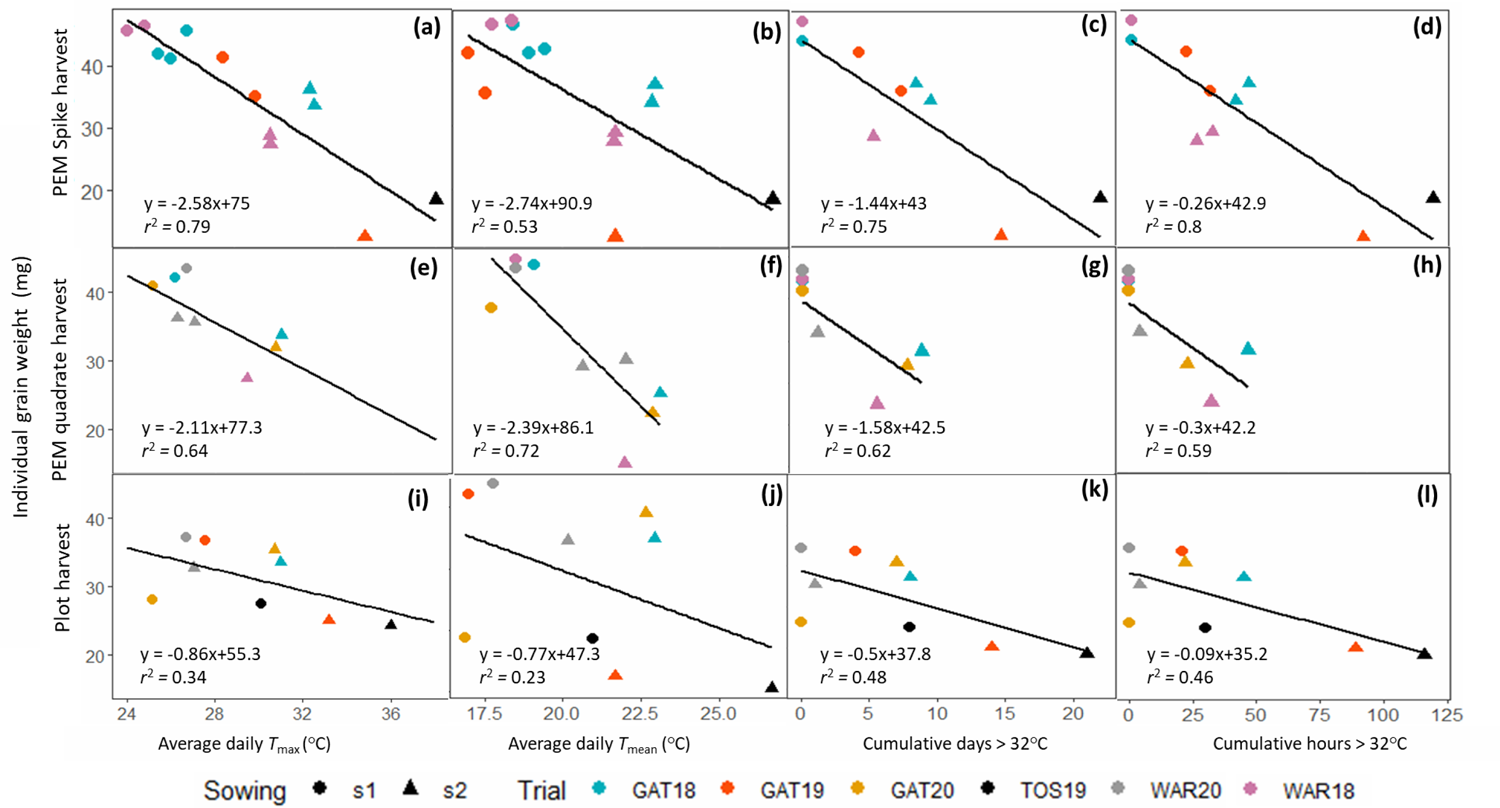
Individual grain weight (IGW) of wheat genotypes in response to post-flowering (a, e, i) daily maximum temperature (*T*max), (b, f, j) mean temperature (*T*mean), cumulative (c, g, k) days and (d, h, l) hours above 32^○^C; all temperature indices were calculated between 0 and 500^○^Cd after flowering. Data correspond to the mean of 20 – 32 genotypes with four independent replicates in each environment. Spike data were collected from the individual spikes (∼20 for each replicate) exposed to heat at synchronised developmental stages. Quadrate data were collected by harvesting 0.5 m linear meter of plants tagged at synchronised flowering in the PEM. Plot data were collected from the whole conventional plots of naturally flowering genotypes. Quadrate and plot data were presented per unit area (m^-2^). Lines were plotted for regressions between mean grain number and temperature indicators that had a regression slope significantly different from zero (P<0.05).

IGW of tested genotypes responded strongly to post-flowering heat. The correlations between IGW and heat post-flowering were closest, with the highest coefficients of determination, for the PEM spike harvests (*r*^2^ = 0.53-0.80), followed by the PEM quadrate harvests (*r*^2^ = 0.47-0.64) and the plot harvests (*r*^2^ = 0.33-0.48; Fig. 2). Compared with averaged IGW under little if any heat stress (HET1), IGW was reduced by an average 3.3%, 3.7% and 1.5% for each hot day (> 32^○^C) during 0-500^○^Cd after flowering for PEM spike (*r*^2^ = 0.75), PEM quadrate (*r*^2^ = 0.62) and plot (*r*^2^ = 0.48) harvests, respectively (Fig. 2 c, g, k). Similarly, for each hot hour (> 32^○^C), IGW was reduced by 0.26, 0.3 and 0.09 mg for PEM spike, the PEM quadrate and the plot harvests, respectively (Fig. 2 d, h, l).

For each 1^○^C increase in average daily *T*_max_ and *T*_mean_ during grain filling, the averaged IGW of the studied genotypes decreased by 2.58 and 2.74 mg, respectively, for PEM spikes (Fig. 2 a, b), and by 2.11 mg and 2.39 mg for PEM quadrates (Fig. 2 e, f).

However, for plot harvests, the correlations between IGW and heat indicators were not as close as for other harvest methods, suggesting that this method is less precise in estimating the IGW loss. For example, *r*^2^ was less than 0.35 for the correlations between IGW of plot harvest and both *T*_max_ and *T*_mean_ (Fig. 2 i, j), while *r*^2^ was above 0.53 when looking those correlations for spike or quadrate PEM harvests (Fig. 2 a-d).

Heat response of total grain weight (‘yield’) was also strongly associated with the studied heat indicators, particularly for PEM spikes (Fig. 3). Each 1^○^C increase in either daily *T*_max_ or *T*_mean_ resulted in a 6.2% reduction in total spike grain weight compared with the minimally-stressed HET1 plants (Fig. 3 a, b). The yield of PEM quadrates (398 g m^-2^) was similar to the yield of conventional plots (368 g m^-2^) under low-stress environments (HET1). However, yield responses to post-flowering temperatures were stronger in plots (*r*^2^ = 0.39-0.65) than in PEM quadrates (*r*^2^ = 0.17-0.24). This was partly due to quadrate not being taken in severe HET3 environments (GAT19-s2 and TOS19-s2). In plots, each 1^○^C increase in average daily *T*_max_ or *T*_mean_ between 0 and 500°Cd post-flowering resulted in 27.8 or 24.7 g m^-2^ yield loss, respectively (Fig. 3i, j). Similarly, for each post-flowering hot day and hour with a temperature above 32^○^C, the yield was reduced by 14.7 and 2.44 g m^-2^, respectively, in plots (Fig. 3 k, l).

**Fig. 3.**
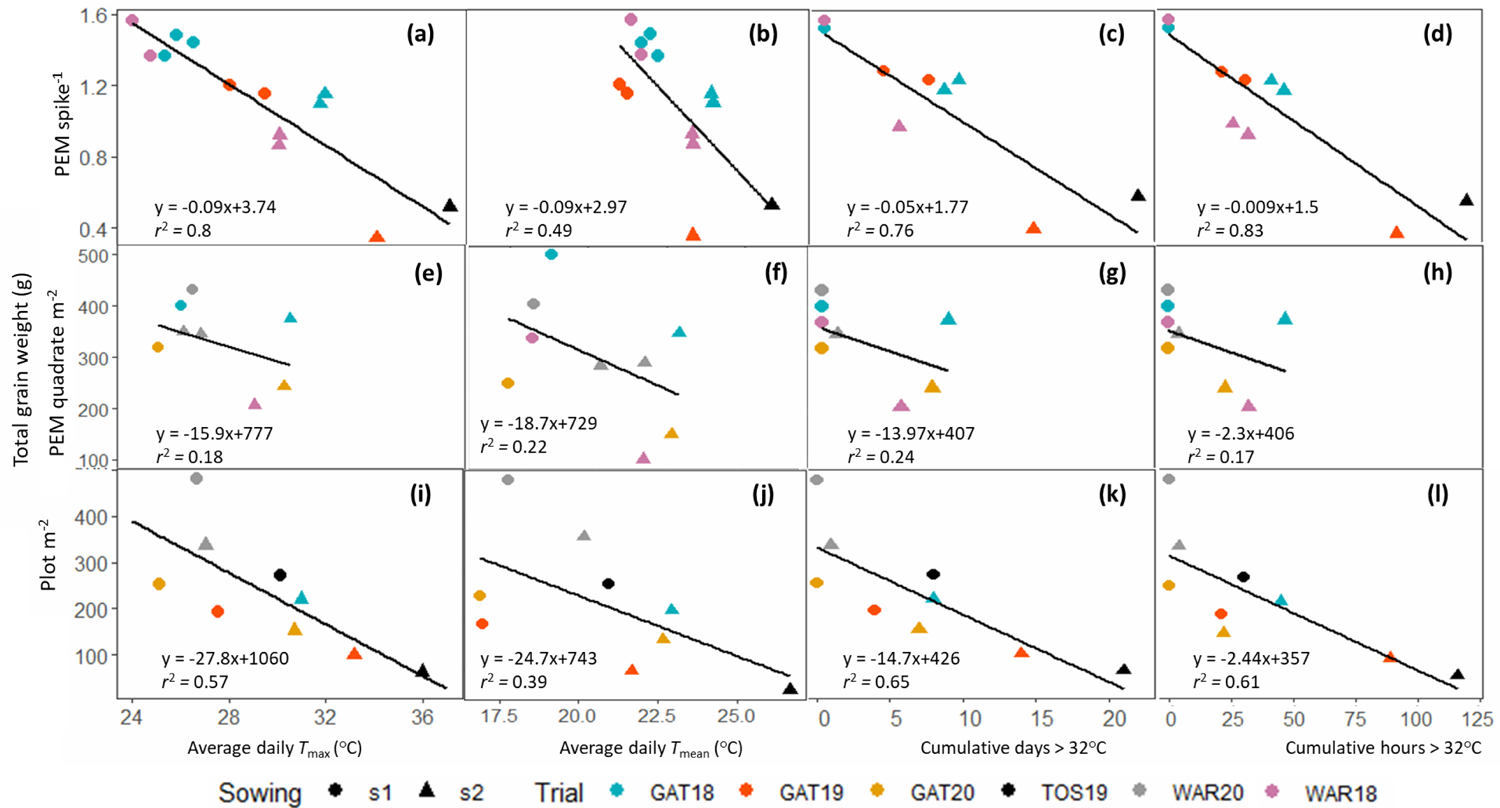
Total grain weight of wheat genotypes response to post-flowering (a, e, i) daily maximum temperature (*T*max), (b, f, j) mean temperature (*T*mean), cumulative (c, g, k) days and (d, h, l) hours above 32^○^C; all temperature indices were calculated between 0 and 500^○^Cd after flowering. Data correspond to the mean of 20 – 32 genotypes with four independent replicates in each environment. Spike data were collected from the individual spikes (∼20 for each replicate) exposed to heat at synchronised developmental stages. Quadrate data were collected by harvesting 0.5 m linear meter of plants tagged at synchronised flowering in the PEM. Plot data were collected from the whole conventional plots of naturally flowering genotypes. Quadrate and plot data were presented per unit area (m^-2^). Lines were plotted for regressions between mean grain number and temperature indicators that had a regression slope significantly different from zero (P<0.05).

### 3.4. Influence of temperature thresholds on the heat response of grain numbers

Reductions in grain yield components (averaged across the tested genotypes) were calculated for each cumulative hot day or hour above different threshold temperatures (from 26^○^C to 32^○^C) during different developmental periods to identify the threshold temperature and duration that allowed best estimation of heat impacts in the tested environments. The computed changes in grain yield components were derived from the individual response curves for different threshold temperatures from 26^○^C to 35^○^C (Fig. 4 a-f). The graph shows how the estimated reductions in grain yield components (for each hot day or hour) vary by increasing the threshold temperatures. The associated coefficients of determination (*r*^2^) are also presented to reflect on the strength of responses to different threshold temperatures (Fig. 4 g-l). Irrespective of the harvest type, the impact of heat on the magnitude of the responses (negative slope) for grain number tended to progressively increase with the increasing temperature threshold, particularly for hot hours (Fig. 4 a, b). However, the coefficient of determination (*r*^2^) tended to either remain relatively unchanged or decline beyond 28^○^C (Fig. 4 g, h). This is most likely because temperatures rarely exceeded 30^○^C early in the season (Fig. S2) and so there were fewer data points for these regressions. The results from the tested conditions suggest that the best estimation for grain number loss is around a threshold temperature of 28^○^C.

**Fig. 4.**
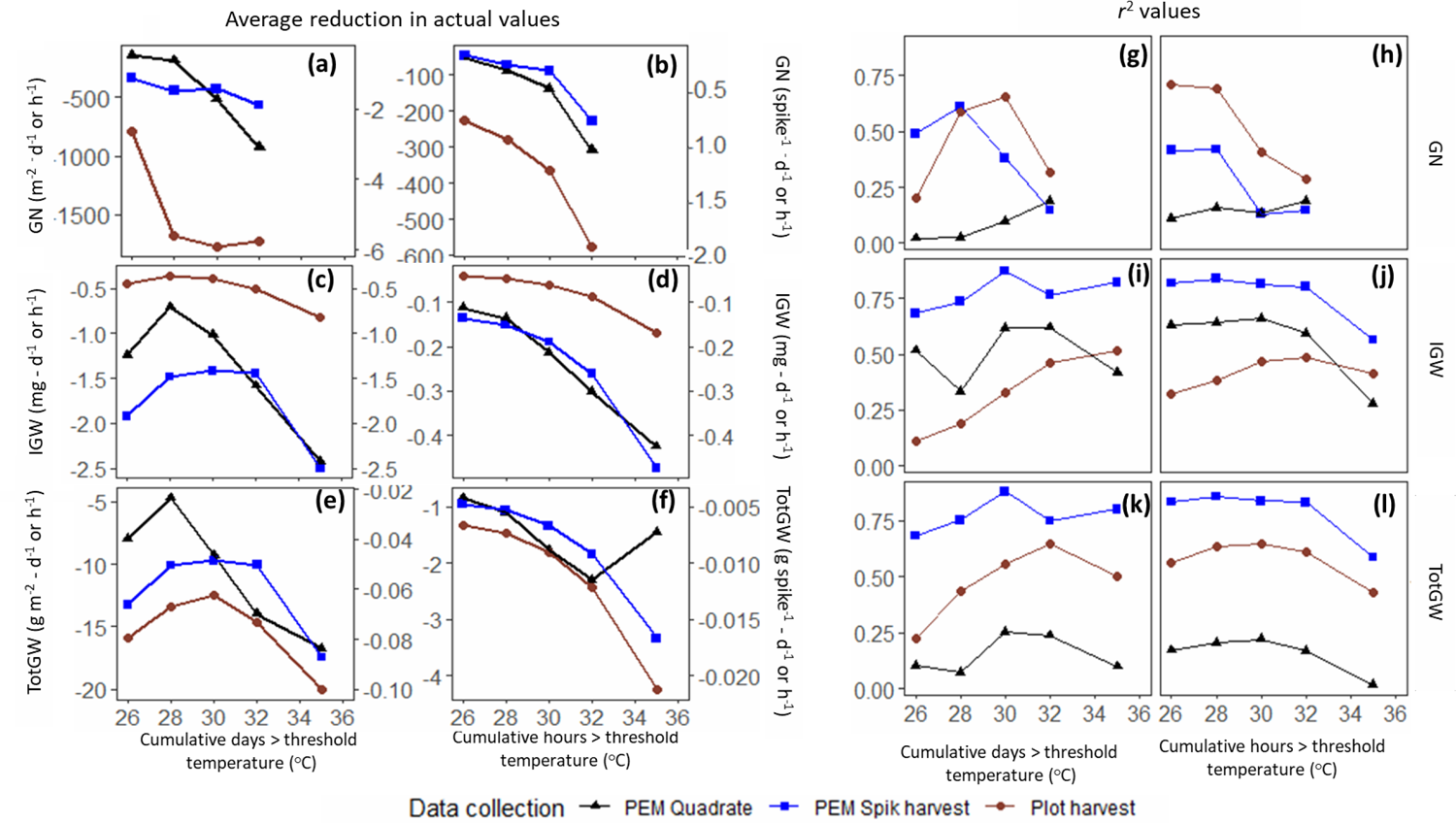
Relative changes in grain yield and its components in response to increasing heat intensity (a to f), and their associated coefficients of determination (g to l). The slope (a-f) and coefficient of determination (g-l) for changes in grain number (GN; a, b), individual grain weight (IGW; c, d) and total grain weight (TotGW; e, f) in response to days (a, c, e) and hours (b, d, f) above threshold temperatures were derived from the individual response curves of different threshold temperatures from 26^○^C to 32^○^C to quantify the reduction for each additional hot day or hour above threshold temperatures on average for 20-32genotypes with four independent replicates. The strange increase in the total grain weight loss found for the hourly increase in threshold temperatures from 32^○^C to 35^○^C in the plot trial (f) resulted from a highly variable genotypic responses (Fig. 4 e) due to variation in the grain filling duration. In (a-f), the right and left-hand Y-axis labels indicate values per unit area (m^-2^) and per spike, respectively. Spike data were collected from the individual spikes (∼20 for each replicate) exposed to heat at synchronised developmental stages. Quadrate data were collected by harvesting 0.5 m linear meter of plants tagged at synchronised flowering in the PEM. Plot data are collected from the whole conventional plots of naturally flowering genotypes.

For conventional plots, the slope of grain number responded sharply to hot hours (more than doubling in absolute value) as the threshold temperature increased from 26^○^C to 28^○^C, but it did not reduce further for 30^○^C or 32^○^C (Fig 4 a). In contrast, the slope of the regression for grain number progressively decreased for the number of hot hours with the increasing threshold temperatures (Fig. 4 b). For example, with each 1^○^C increase in threshold temperature from 26^○^C to 32^○^C, wheat genotypes lost an additional 57 grains m^-2^ on average in conventional plots (Fig. 4 b). This suggests that using hot hours rather than hot days for estimating grain number loss was more responsive, particularly at the plot level during this developmental period.

For grain number per spike collected on stems at synchronised development stages (PEM), the slope of grain number response to hot hours decreased mainly when the temperature threshold for hot hours was increased from 30^○^C to 32^○^C (Fig. 4 b). In other words, in the tested conditions, the grain number of PEM spikes was less responsive to increases in pre-flowering threshold temperatures below 30^○^C. However, coefficients of determination (r^2^) were the strongest for low threshold temperature of 28-30°C (Fig. 4 h) as higher temperatures occurred pre flowering in only a few environments (Fig. S2).

For PEM quadrates, loss in grain number progressively increased with the increasing threshold temperatures both for hot hours and hot days (Fig. 4 e, f), but the coefficients of determination were low (*r^2^* < 0.2) (Fig. 4 g, h), likely due to only a few heat events before flowering in the tested environments with PEM quadrates resulting in a limited number of data points for the regression curves (Fig. S2).

### 3.5. Influence of the temperature thresholds on the heat response of grain weight

Loss in IGW and total grain weight strongly responded to increasing post-flowering (0-500^○^Cd) threshold temperatures (Fig. 4 i-l). Across the tested temperature thresholds for days and hours, these responses were strongest for PEM spikes, followed by PEM quadrates and plots (Fig. 4 i, k). For instance, for temperature threshold for days, *r*^2^ for IGW was maximum at 0.88 for PEM spikes, 0.62 for PEM quadrates and 0.36 for conventional plots (Fig. 4 i); for total grain weight r^2^ was maximum at 0.87 for PEM spike, 0.24 for PEM quadrate, and 0.64 for conventional plots (Fig. 4 k). Further, for conventional plots, the response of IGW and total grain weight to temperature threshold became progressively stronger as *r*^2^ increased from 28^○^C to 32^○^C, particularly for hot days and then slightly declined for 35^○^C (Fig. 4 i-l). Hence, for the tested conditions in conventional plots, estimating heat impacts on IGW and grain yield appeared most precise when considering highly stressful conditions (i.e. high-temperature threshold of 32^○^C) that occur relatively frequently (Fig. S3). In contrast, it is less precise at temperatures such as 35°C, which is relatively rare in tested environments. Additionally, the impact could typically be more precisely quantified using hot hours above threshold temperature rather than hot days (Fig. 4 i-l, S3).

As expected, the heat impact per hot day or hour on either IGW or total grain weight generally increased when considering greater temperature thresholds (Fig. 4 c-f). However, this was not always the case for non-stressful or less stressful conditions when considering daily data (Fig. 4 c, e). In contrast, when considering hourly data (Fig. 4 d, f) the impact of heat on IGW and total grain weight (weight reduction hour^-1^) progressively intensified as threshold temperature increased from 26°C to 35°C, except for the total grain weight of PEM quadrate under 35^○^C. On average, for each 1^○^C increase in hourly temperature threshold (from 26°C to 35°C), wheat genotypes experienced an additional reduction in IGW of 0.04 mg per hot hour for both PEM spikes and quadrates (Fig. 4 d) and a yield reduction of 0.32 g m^-2^ h^-1^ for conventional plots (Fig. 4f).

### 3.6. Impact of the timing of post-flowering heat

The impact of post-flowering heat on grain weight was further investigated by exploring how the timing of the first heat event affects IGW. Based on the data presented above, and as previously suggested by Collins et al. (2000), a heat event was defined as at least a 4 h cumulative period of temperatures above 32^○^C in order to identify the first heat event. In the tested conditions, IGW was not or only little affected when intense heat only started occurring during the late period of grain development, e.g. 400°C or 500^○^Cd after flowering. However, IGW responded strongly to the timing of the first heat event when it occurred early after flowering (Fig. 5).

**Fig. 5.**
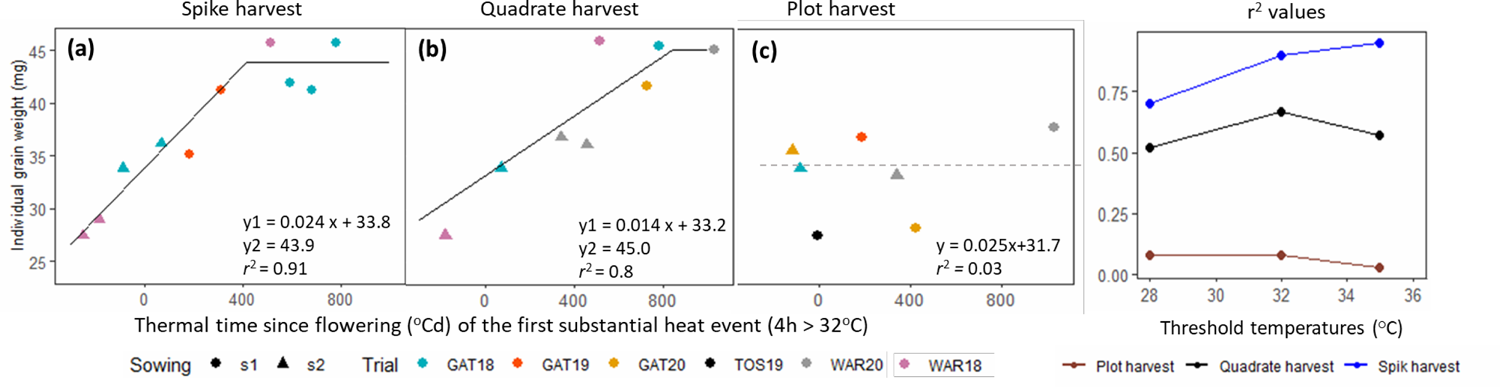
Individual grain weight of wheat genotypes plotted against the timing relative to the flowering of the first heat event (4 cumulative hours > 32^○^C) for (a) spikes with synchronised phenology, (b) quadrates with synchronised phenology, and (c) conventional plots (without synchronised phenology), together with (d) associated coefficients of determination for different temperature thresholds (first heat event being defined as 4 hours > *Tthres*). Spike data were collected from individual spikes (∼20 for each replicate) exposed to heat at synchronised developmental stages in the PEM. Quadrate data were collected by harvesting 0.5 m linear meter of plants tagged at synchronised flowering in the PEM. Plot data were collected from the whole conventional plots of naturally flowering genotypes. Data are the mean of 20 – 32 genotypes with four independent replicates. Slope and *r*^2^ values are presented for trials from HET1 and HET2 only.

For PEM spikes and quadrates, wheat genotypes were compared at synchronised development stages. The timing of the first heat shock affected genotypes at a similar stage relative to flowering in the PEM, but not in conventional field plots. Before late grain filling in HET1-2 environments, for each 100^○^Cd delay (∼5 days) in the start of extreme heat shocks (4h > 32°C), the tested genotypes produced on average 2.4 mg (*r*^2^ = 0.91) and 1.4 mg (*r*^2^ = 0.8) larger grains at the spike and quadrate levels in the PEM, respectively (Fig. 5 a, b). By contrast, no clear response to the timing of the first heat event was observed for IGW in conventional plots (Fig. 5 c), possibly because of phenological variation across the tested genotypes when natural heat events occurred.

The importance of the timing of heat stress on grain filling is evident when comparing GAT18-s2 and WAR19s2 (Fig. 6 a-c). The plants in GAT18-s2 produced bigger grains than WAR18-s2 despite being subjected to more hot hours (T > 28^○^C) during the grain filling period (0-500°Cd after flowering) (Fig. 6 a, c). By contrast, when focusing solely on the first 100°Cd post-flowering, the plants responded similarly to the cumulative hot hours in both trials (Fig. 6 b). Crops in WAR18-s2 received twice as many hours above 28^○^C compared to GAT18-s2 crops; and they experienced significantly more reduction in IGW at maturity, despite receiving overall fewer post-flowering hot hours (76 h in WAR18-s2 compared with 108 h in GAT18-s2 between 0-500^○^Cd after flowering; Fig. 6). This example illustrates the complexity of working with fluctuating natural environments, while it also highlights that heat events occurring during the early grain development are more impactful than those occurring later.

**Fig. 6.**
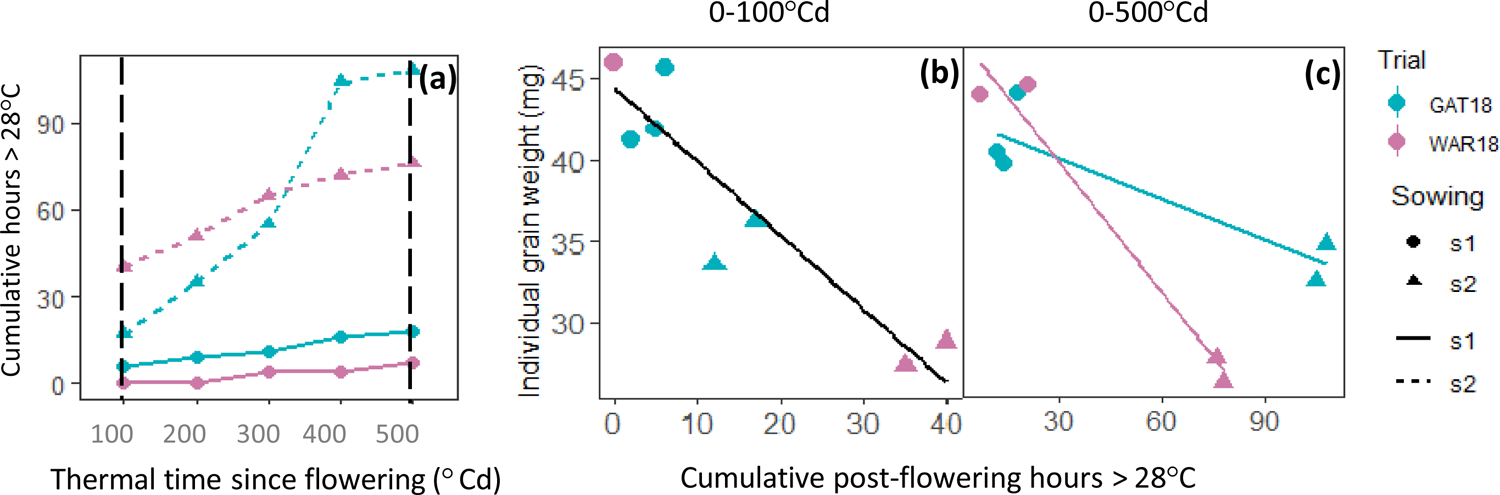
Illustration of the importance of the timing of heat events on individual grain weight (IGW; b-c) in two trials with contrasting patterns of heat events (a). (a) Cumulative hours > 28^○^C for different post-flowering periods (0-100^○^Cd, 0-200^○^Cd, 0-300^○^Cd, 0-400^○^Cd and 0-500^○^Cd after flowering) for the first tagging of each of two 2018 trials. The number of hours > 28^○^C during the first 100^○^Cd after flowering (black dashed vertical line in (a)) highlights warmer conditions in early grain filling in WAR18-s2 compared to GAT18-s2, while this environment was cooler towards the end of the grain filling. Changes in IGW in response to cumulative hours > 28^○^C were consistent across environments during 0-100^○^Cd after flowering (b). However, IGW was greater in GAT18-s2 than WAR18-s2 despite warmer average grain-filling temperature (0-500^○^Cd after flowering; c). This was due to warmer temperatures in GAT18-s2 occurring late in the grain filling period (400^○^Cd to 500^○^Cd), when IGW was no longer highly sensitive to heat. In (b) and (c), IGW is presented for all taggings of 2018 trials. Data were collected from individual spikes (∼20 for each replicate), and averaged for all 32 studied genotypes and replicates.

## 4. Discussion

### 4.1. High impact on grain number and grain weight is highly stage sensitive

Grain number was strongly correlated with high temperatures between 200-300^○^Cd (∼10-15 days) before flowering (Fig. 1). This period corresponds to pollen meiosis – the most stress-sensitive phase of pollen development (Dolferus et al., 2013; Masoomi-Aladizgeh et al., 2021). Damage to microspores during this phase is irreversible, translating into grain number loss at maturity, irrespective of temperature later during development (Dolferus, 2014; Ji et al., 2010). While field-based studies have mainly focused on heat impact in response to the whole pre-flowering period (Thistlethwaite et al., 2020; Talukder et al., 2014), controlled environment studies also linked the grain set in wheat with high temperatures during pollen meiosis (10-15 days before flowering) or around anthesis (Prasad and Djanaguiraman, 2014; Saini and Aspinall, 1982). However, the impact of heat on pollen viability has previously been studied only *in-vitro* (Impe et al., 2020) or in controlled environment under extremely high temperatures (i.e. 35°C, Prasad and Djanaguiraman, 2014). For example, three hot days (35/22°C day/night) during meiosis reduced the pollen viability of wheat genotypes by 57% (Bokshi et al., 2021) and grain number per spike by 9% (Thistlethwaite et al., 2020). Interestingly, no significant correlations between grain number and heat indicators between 100°Cd before to 100°Cd after flowering were found for PEM harvests, suggesting that grain number is more susceptible to mild heat (∼28°C) during the early pollen developmental phase than around anthesis.

IGW of the tested genotypes responded strongly to high temperatures from 0-500^○^Cd post-flowering. In PEM harvests, for each 1^○^C increase in post-flowering daily *T*_max_, IGW was reduced by 2.6 mg (spike harvest) and 2.1 mg (quadrate harvest) compared (Fig. 2 a, e). The high sensitivity of IGW to post-flowering heat has already been reported for plants grown in controlled environments (Stone and Nicolas, 1995; Tashiro and Wardlaw, 1990). Similar to grain number, previous field-based studies tend to report the impact of heat for the total post-flowering duration rather than for a short period specific to a given development stage (Telfer et al., 2018; Thistlethwaite et al., 2015). In contrast, synchronised phenology under PEM and detailed tracking of heat events (timing and intensity) in the current study allowed precise quantification of heat-induced IGW loss in natural field conditions.

That the response of wheat to heat is strongly influenced by developmental phase was also evident from a close correlation (*r*^2^ = 0.91) between IGW and the timing of the first post-flowering extreme heat event (4h > 32^○^C) for spikes with synchronised development (Fig. 5 a). High temperature during early grain filling caused a maximum reduction in IGW, but grains become progressively less sensitive to heat at later stages (e.g. > 400^○^Cd post-flowering; Fig. 6). This is in adequation with the findings of Stone and Nicolas (1995), who found lower sensitivity of wheat grains to heat during later phases of development (30 days after anthesis) than 15-20 days after anthesis. The grain development period can be divided into three different phases: (phase I) rapid cell proliferation, (phase II) grain filling, and (phase III) maturation (Ellis, 1999). Our study suggested that IGW of tested wheat genotypes is more sensitive to heat during early phases of grain development (i.e. I and II) compared to the maturation phase (III). We further compared the impact of high temperature during the first 100^○^Cd (cell proliferation phase) and most of the grain filling duration (0-500^○^Cd after flowering). Results suggested that intensity and duration of heat during phase I (∼0-10 days after flowering), is a more critical determinant of final grain size than total post-flowering heat (Fig. 6). This suggests that high temperature during early grain development can irreversibly impair cellular division and enlargement (Girousse et al., 2021), impacting the final grain size irrespective of temperature at later stages, as observed in the current study (Fig. 6).

### 4.2. Which heat indicators to characterise heat stress impacts?

Crops responses to heat stress are complex as they depend on the timing, intensity, duration of the stress as well as other factors such as acclimation, making them hard to model quantitatively. Response of yield components to heat in this study depended heavily on the temperature indicator considered such as *T*_max_, *T*_mean_ or cumulative days or hours > threshold temperatures. For example, the coefficient of determination (*r*^2^) between grain number and each of the studied temperature indicators ranged from 0.18 to 0.61 for PEM spikes (Fig. 1 a-d). This highlights the importance of selecting a suitable temperature indicator to quantify heat impacts with greatest sensitivity and accuracy.

Heat-induced losses in grain number vary widely across published studies, likely due in part to the differences in growing conditions, heat indicators considered and genetic backgrounds. Even in controlled environment, impacts of heat on grain number may vary. For instance, three hot days (35°C) around anthesis was reported to reduce grain number by 22% (Thistlethwaite et al., 2015), while in another study, a single day hot day (T > 35°C) reduced grain number by 24% (Talukder et al., 2014). In the current study, we used various heat indicators in an attempt to better understand how the estimated grain loss varies with varying heat intensities. We found that grain number responded strongly to pre-flowering heat indicators in spike PEM and conventional plots (Fig. 1 i-l). For instance, by increasing daily *T*_max_ from 21.5 to 32°C (in average between 200 and 300°Cd before flowering), grain number was impacted by 7% for each 1°C in conventional plots (Fig. 1 i). However, understanding response to increased temperature is complex, and impact due increased average temperature (thermal time) and acceleration of phenology were also observed. In conventional plots, for each 1°C increase in *T*_mean_ during the 200-300°Cd pre-flowering period, the tested genotypes produced 10% fewer grains in harvests with none or little stressed HET1 controls (Fig. 1 j). This reduction is higher than reported by Fischer (1985), who observed that a 1°C increase in daily mean temperature during the spike growth period (∼30 days prior to flowering) reduces grain numbers by 4%, at least partly due to differences in growing conditions (e.g. lower average temperature varying between 14 and 22°C). Our study shows that with increasing threshold temperature, in particular for the definition of cumulative ‘hot’ hours, grain loss significantly decreases (Fig. 4 b, d). For each hot hour (> 28°C) between 300 and 200°Cd before flowering, grain number was estimated to decrease by 2.5% in conventional plots (Fig. 1 l).

In the current study, changes in grain number for spikes with synchronised flowering were more strongly associated with pre-flowering *T*_max_ than pre-flowering *T*_mean_, suggesting an impact of heat stress (and not just a response to thermal time) on grain loss (Fig. 1 a, b). This is in agreement with the report of Ferris et al. (1998), who found that grain number in wheat responded strongly to *T*_max_ but not to *T*_mean_ when the crop was exposed to heat during flowering. Similarly, Musa et al. (2021) also suggested *T*_max_ and the number of hot days as the best indicators of grain number loss in wheat genotypes under hot environments (*T*_max_ > 32°C). However, in the absence of extreme heat, e.g. only few days of *T*_max_ > 32°C, grain number may respond more strongly to *T*_mean_ (He et al., 2020, Ye et al., 2021). In conventional plots (without synchronised flowering), the response of grain number was relatively stable across all studied temperature factors (*r*^2^ between 0.55 and 0.69; Fig. 1 i-l).

IGW also responded more strongly to post-flowering *T*_max_ than *T*_mean_ (0-500°Cd after flowering), with *r*^2^ (averaged across harvest types) of 0.59 and 0.44 for *T*_max_ and *T*_mean_, respectively (Fig. 2). Compared with grain number, the impact of heat on IGW was estimated for a broader window (0-500°Cd after flowering). For each 1°C increase in post-flowering *T*_max_ and *T*_mean_, IGW was reduced by 2.58 and 2.74 mg, respectively, for spikes with synchronised development (Fig. 2 a, b). A reduction of 2.80 mg in IGW was also previously reported for each 1°C increase in *T*_mean_ during the grain-filling phase of irrigated wheat by Wiegand and Cuellar (1981). As expected, the estimated impact of *T*_max_ on IGW was smaller and not as clear for PEM quadrates and conventional plots compared to PEM spikes, e.g. 2.11 mg loss per degree in *T*_max_ for PEM quadrate (*r*^2^ = 0.64), and 0.86 mg loss per degree in *T*_max_ for conventional plot (*r*^2^ = 0.34).

In regard to yield, a loss of 14.7 g m^-2^ (*r*^2^ = 0.56) was found for each hot post-flowering day > 32°C in conventional plots (Fig. 3 k), which is similar to 16.1 g m^-2^ for each hot day (> 30°C) between 100 and 600°Cd post-anthesis under field conditions (Telfer et al., 2018). Controlled environment studies suggested a 23% reduction in wheat grain yield after 4 post-flowering hot days (> 35°C) (Stone and Nicolas, 1994), which is slightly lower than our estimation, i.e. 4.6% (*r*^2^ = 0.5) for each hot day > 35°C compared to HET1 environments in conventional plots (Fig. 4 e). Similarly, Thistlethwaite et al. (2020) reported a 40 g m^-2^ grain yield reduction for each 1°C increase in *T*_mean_ under irrigated field conditions. However, this value is substantially higher than our estimated grain yield loss of 24.7 g m^2^ in conventional plots (Fig. 3 j). Across the literature, a broad range of yield loss (3 – 18%) has been reported for each 1°C rise in post-flowering atmospheric temperature above the optimum in well-watered wheat crops (He et al., 2019; Mondal et al., 2013; Ullah, 2018). These variations may be explained, at least partially, by the difference in growing environments (e.g., number of hot days and warm nights, other environmental factors also influencing crop growth and development), experimental methodologies (e.g. period considered to calculate the mean temperature) and studied genotypes.

In the current study, IGW and total grain weight loss were estimated more reliably with hours rather than days of cumulated heat, particularly for some of the low and high examined temperature thresholds (e.g. 26 and 35^○^C, Fig. 4 c-f).

### 4.3. Synchronised phenology across genotypes improved estimation of heat stress impact on individual grain weight

The impact of heat stress was compared for spikes at synchronised development stage, quadrates at synchronised development stage and conventional plots at a range of developmental stages (not synchronised). In this study, IGW was more reliably estimated with the synchronised phenology than in conventional plots, especially when considering individual spikes that were at synchronised development stage. Coefficients of determination (*r*^2^) from regressions between IGW and heat indicators (averaged across all tested indicators) were 0.72, 0.58 and 0.40 for PEM spikes, PEM quadrate and conventional plots, respectively (Fig. 2). For instance, for each 1^○^C increase in *T*_max_ or each hot day (> 32^○^C) during 0-500^○^Cd after flowering, IGW of the tested genotypes reduced by 6% (*T*_max_) or 3.3% (*T* > 32^○^C) for PEM spike and 5% (*T*_max_) or 3.7% (*T* > 32^○^C) for PEM quadrate and 2.6% (*T*_max_) or 1.5% (*T* > 32^○^C) for conventional plots, respectively. For spike harvest, tillers of synchronised phenology experienced identical heat intensity during similar developmental stages, so that IGW responses to heat indicators were stronger, with both a greater estimated impact of heat and greater *r^2^*, in PEM spikes than in PEM quadrates, and even more so compared to conventional plots (Fig. 2 and 4).

In conventional plots with unsynchronised flowering (due to differences in genotype maturity types and asynchronism between main stems, primary and secondary tillers), the tested genotypes/spikes experienced heat events at different developmental phases, which weakened IGW response to heat indicators (Fig. 2 and 4). Different responses of IGW from different tillers to heat stress around flowering have already been confirmed under controlled environments (Aiqing et al., 2018), in particular, due to their asynchronism (Chenu and Oudin, 2019). Highly development-phase-specific impact of heat on IGW was recorded in the current study (Fig. 5 and 6). For spikes with synchronised phenology, IGW responded strongly (*r*^2^ = 0.95) to the timing of the first heat event (4 hours > 35^○^C), but for conventional plots, no significant correlation was observed (*r*^2^ = 0.03). This suggests that quantification of IGW loss in response to heat could be achieved more reliably by focusing on the spikes of synchronised phenology and, to a lesser extent, focusing on whole plants in quadrates of synchronised phenology (PEM quadrates; Fig. 5).

Further, total grain weight (spike^-1^ or m^-2^) was reliably estimated from both spike and plot harvests (Fig. 3), and relatively stronger correlations between heat indicators and grain weight were observed at spike than at plot level. For instance, for each 1^○^C increase in *T*_max_ during post-flowering 0-500^○^Cd, total grain weight was reduced by 6.2% (*r*^2^= 0.8) and 6.4% (*r*^2^= 0.57) for spikes and plots, respectively, compared with plants from optimum HET1 environments. Similarly, correlations for grain yield, with hot hours or hot days for spike harvest, were stronger than for the conventional plots across any tested temperature (Fig. 4 k-l).

The current study suggests that heat impact on wheat grain yield can be most accurately estimated by focusing on specific developmental phases. For example, grain number responded strongly to heat indicators for a short period of ∼ 5 days (200-300^○^Cd before flowering), presumably during the critical period of pollen meiosis (Dolferus et al., 2013; Masoomi-Aladizgeh et al., 2021). Although IGW correlated strongly to the heat indicator for a broader range time (0-500^○^Cd after flowering), grain weight loss was highly development-phase-specific with significantly more sensitivity to heat during the early development phase (Fig. 6). Changes in yield components were also more accurately quantified using heat indicators such as *T*_max_ and the number of hot hours on wheat plants with synchronised phenology than *T*_mean_ and hot days.

The strongest relationships for heat responses of grain number were found for a threshold temperature of 28°C from 200 to 300^○^Cd before flowering. For IGW, best-suited thresholds were >= 28°C when considering cumulated hot hours, or thresholds were >= 32°C for cumulated hot days.

## 5. Conclusions

The best estimations for yield-component response to heat were achieved with spikes or quadrates at synchronised development. In the tested environments, the strongest relationships for responses of grain number were found for average daily maximum temperature and cumulative hours with the temperature above 28°C during the period from 200 to 300^○^Cd before flowering. IGW responded most strongly to maximum temperature and cumulative hours with the temperature above 28°C during the early to mid grain filling stages (0-500°Cd after flowering).

Our study suggested timing of the first heat event during early grain filling (0-100°Cd post flowering) is critical for final grain size. Results indicated that the impact of natural heat on wheat grain yield components in the field could be accurately estimated with the selection of appropriate experimental techniques and heat indicators. Optimising field experimental techniques will be particularly important for screening the relative heat tolerance of genotypes in breeding programs.

## Acknowledgments

The research was made possible thanks to the support of The University of Queensland and the Queensland Government, Department of Agriculture and Fisheries. Najeeb Ullah was supported by a Queensland Government as an Advance Queensland Fellowship, and by The University of Queensland with an Amplify Fellowship. We acknowledge the assistance of Ian Broad (Department of Agriculture and Fisheries Queensland) with operating of weather stations across the studied sites and Thaís Helena Godoy Sanches for sample processing and assistance in data collection.

## Abbreviations

*T*_max_: average daily maximum temperature

*T*_mean_: average daily mean temperature

*T*_thresh_: threshold temperature

IGW: individual grain weight

PEM: photoperiod extension method

GN: grain number

TotGW: total grain weight

tSow1: time of sowing1

tSow2: time of sowing 2

GAT: Gatton

TOS: Tosari

WAR: Warwick

*Tair_h_*: hourly air temperature

VPD: vapour pressure deficit

