## Supplementray data for "Pre- and post-flowering impacts of natural heat events on yield components in wheat"

**Table S1**: Characteristics of wheat genotypes used in the study

| Genotypes | Type | Pedigree | Characteristics | Reference | Year tested |
|---|---|---|---|---|---|
| Berkut | Cultivar | Irena/Baviacora-M-92//Pastor | Heat tolerance | (Thistlethwaite et al., 2020) | 2018, 2019, 2020 |
| Corack | Cultivar | MACHETE/84W129-504*2/5/VICAM S 71//CIANO F 67/SIETE CERROS/ 3/KAL/BB/4/TM56 | Australian Premium White (APW) variety | (GRDC, 2018) | 2018, 2019 |
| Dharwar Dry | Cultivar | DWR39/C306//HD2189 | Drought tolerant, stay-green phenotype | (Manschadi et al., 2008) | 2018, 2019 |
| Drysdale | Cultivar | Hartog*3/Quarrion | Superior transpiration efficiency | (Fletcher et al., 2018; Lobell et al., 2015; Richard et al., 2015) | 2018, 2019, 2020 |
| EGA Gregory | Cultivar | Pelsart/2*Batavia | Long-season elite cultivar with broad adaptation | | 2018, 2019, 2020 |
| EGA Wylie | Cultivar | INIA F 66 / GAMUT// COOK/3/JUPATECO F 73/TR 59 | Disease resistance | (Zheng et al., 2014) | 2018, 2019, 2020 |
| FAC10-16 | Elite breeding line | 10CB-F/W234 | Disease resistance | (Dinglasan et al., 2016) | 2018, 2019 |
| Fang | Cultivar | ANNUELLO/2*STYLET | Heat tolerance | (Skylas et al., 2002) | 2018 |
| GK ARON/AG | Elite breeding line | GK ARON/AG SECO 7846//2180 /4/2*MILAN/KAUZ//PRINIA/3/BAV92 | Heat tolerance | (Thistlethwaite et al., 2020) | 2018, 2019 |
| GOU/Sokoll | Elite breeding line | GOUBARA-1/2*Sokoll | Heat tolerance | (Thistlethwaite et al., 2020) | 2018, 2019, 2020 |
| Hartog | Cultivar | Vicam 71//Ciano 's'/Siete Cerros/3/Kalyansona/Bluebird | Drought sensitive, senescent phenotype | (Christopher et al., 2008; Richard et al., 2015) | 2018, 2019, 2020 |
| Janz | Cultivar | 3-AG-3/4*CONDOR//COOK | Drought sensitive, senescent phenotype | Christopher et al., 2008 | 2018, 2019, 2020 |
| Mace | Cultivar | Wyalkatchem/Stylet//Wyalkatchem | Benchmark cultivar for yield across southern Australian cropping regions | (GRDC, 2018) | 2018, 2019, 2020 |
| Mace-177 | Elite breeding line | Mace/Seri-M82 | Drought adaptation | (Christopher et al., 2021) | 2018, 2019 |
| PBW343 | Cultivar | NORD-DESPREZ/VG-1944//KALYANSONA//BLUEBIRD/3/YACO(SIB)/4/VEERY-5 | Heat tolerant | (Thistlethwaite et al., 2020) | 2018 |
| RIL114 | Elite breeding line | UQ01484/RSY10//H45 | Pre-harvest sprouting tolerance | (Hickey et al., 2009) | 2018, 2019 |
| SB003 | Elite breeding line | Seri-M82/Babax | Heat sensitive, low stem water soluble carbohydrates | (Dreccer et al., 2009; Olivares-Villegas et al., 2007; Rattey et al., 2009; Ullah and Chenu, 2019) | 2019, 2020 |
| SB062 | Elite breeding line | Seri-M82/Babax | Heat tolerant, high stem water soluble carbohydrates | (Dreccer et al., 2009; Olivares-Villegas et al., 2007; Rattey et al., 2009; Ullah and Chenu, 2019) | 2018, 2019, 2020 |
| Scout | Cultivar | Sunstate/QH71-6//Yitpi | Elite cultivar with broad adaptation | | 2018, 2019, 2020 |
| Scout-136 | Elite breeding line | Scout/RIL114 | Drought sensitive, senescent phenotype | (Christopher et al., 2021) | 2018, 2019, 2020 |
| Seri-M82 | Elite breeding line | Kavkaz/4/Saric F 70///Lerma Rojo 64A/Inia F66//Inia F66/Yecora F70/5/II-26992 | Dense root system, stay-green phenotype | (Christopher et al., 2008; Olivares-Villegas et al., 2007) | 2018, 2019, 2020 |
| Sokoll | Cultivar | Pastor/3/Altar84/AE.SQ (TR.TA)//OPATA-M-85 | Heat tolerant | (Thistlethwaite et al., 2020) | 2018, 2019, 2020 |

| | | | | | |
|---|---|---|---|---|---|
| Sokoll//FRTL | Elite breeding line | SOKOLL//FRTL/2*PIFED | Heat tolerant | (Thistlethwaite et al., 2020) | 2018, 2019, 2020 |
| Spitfire | Cultivar | Drysdale/Kukri | Heat tolerant, high grain protein content | | 2018, 2019 |
| SSrT17 | Elite breeding line | Double backcross for the TIN gene into the free-tillering Silverstar background | Low tillering (*tin1* allele), big grains | (Mitchell et al., 2008) | 2019 |
| SSrW35 | Elite breeding line | Double backcross for the TIN gene into the free-tillering Silverstar background | High tillering (wild type allele) | (Mitchell et al., 2008) | 2019 |
| Suntop | Cultivar | Sunco/2*Pastor//SUN436E | Elite cultivar with broad adaptation; superior transpiration efficient | (Collins et al., 2021; Richard et al., 2015) | 2018, 2019, 2020 |
| Suntop_1 | Elite breeding line | Suntop /Dharwar Dry | Drought tolerant, stay-green phenotype | (Christopher et al., 2021) | 2018, 2019 |
| Suntop _198 | Elite breeding line | Suntop/SB062 | Drought tolerant, stay-green phenotype | (Christopher et al., 2021) | 2018, 2019, 2020 |
| Suntop _52 | Elite breeding line | Suntop /Dharwar Dry | Drought tolerant, stay-green phenotype | (Christopher et al., 2021) | 2018, 2019, 2020 |
| WH 542 | Cultivar | BJY/JUP//URES | Heat tolerant | (Thistlethwaite et al., 2020) | 2018 |
| Yitpi | Cultivar | Condor/Gabo | Long coleoptile, very susceptible to yellow spot | | 2018, 2019, 2020 |
| ZWB10-37 | Elite breeding line | Tacupeto F2001/Brambling//Kiritati | High yield in CIMMYT-Australia-ICARDA Germplasm Evaluation | (Dinglasan et al., 2016) | 2018, 2019, 2020 |
| ZWW10-128 | Elite breeding line | ESDA/KKTS | High yield in CIMMYT-Australia-ICARDA Germplasm Evaluation | (Dinglasan et al., 2016) | 2018, 2019 |
| ZWW10-50 | Elite breeding line | Onix/4/Milan/Kauz//Prinia/3/BAV92 | High yield in the CIMMYT-Australia-ICARDA Germplasm Evaluation | (Dinglasan et al., 2016) | 2018, 2019, 2020 |

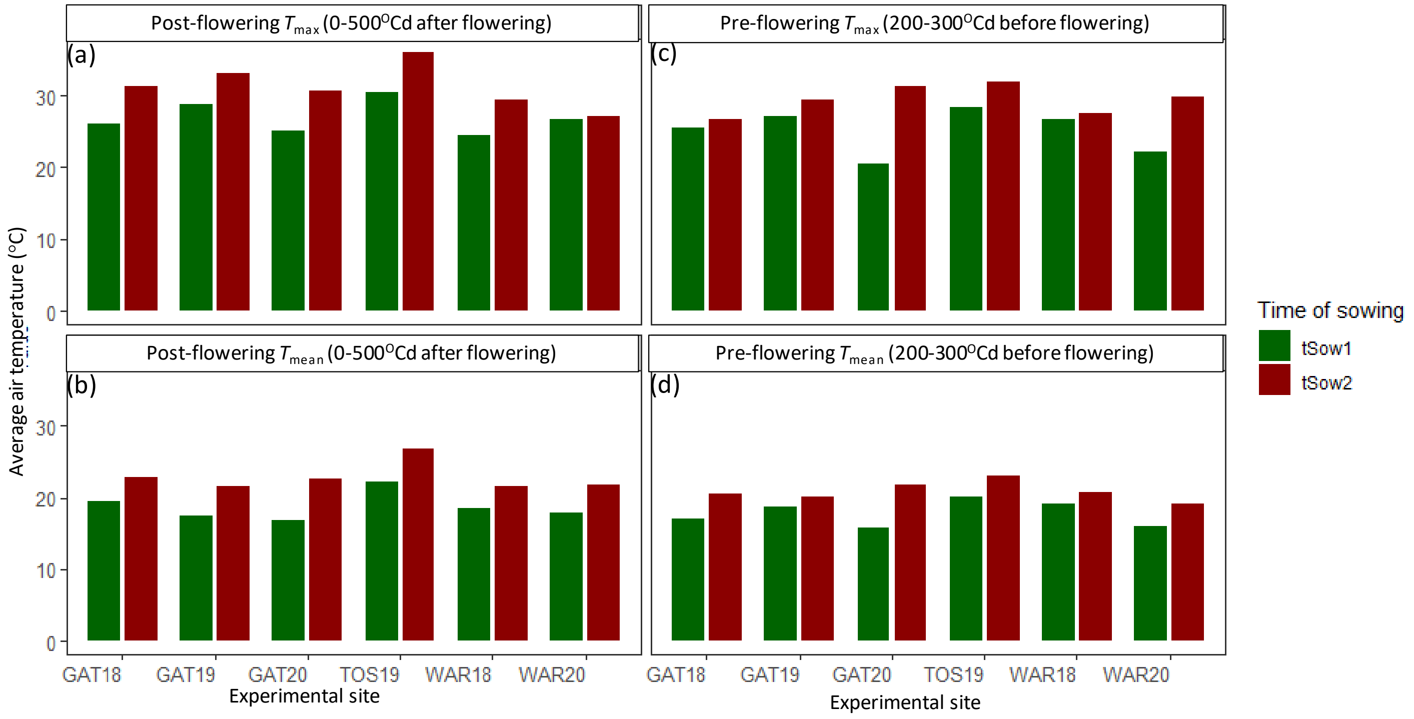


Figure S1: Intensity of heat experienced by wheat crops during sensitive developmental phases before (a) and after (b) flowering across trials and taggings. $T_{max}$ and $T_{mean}$ are average daily maximum and mean temperatures, respectively. T1, 1st cohort of stems tagged at flowering, T2: 2nd cohort of stems tagged at flowering, T3: 3rd cohort of stems tagged at flowering.

Commented [KC1]: In a, xlabel: from 300 to 200oC

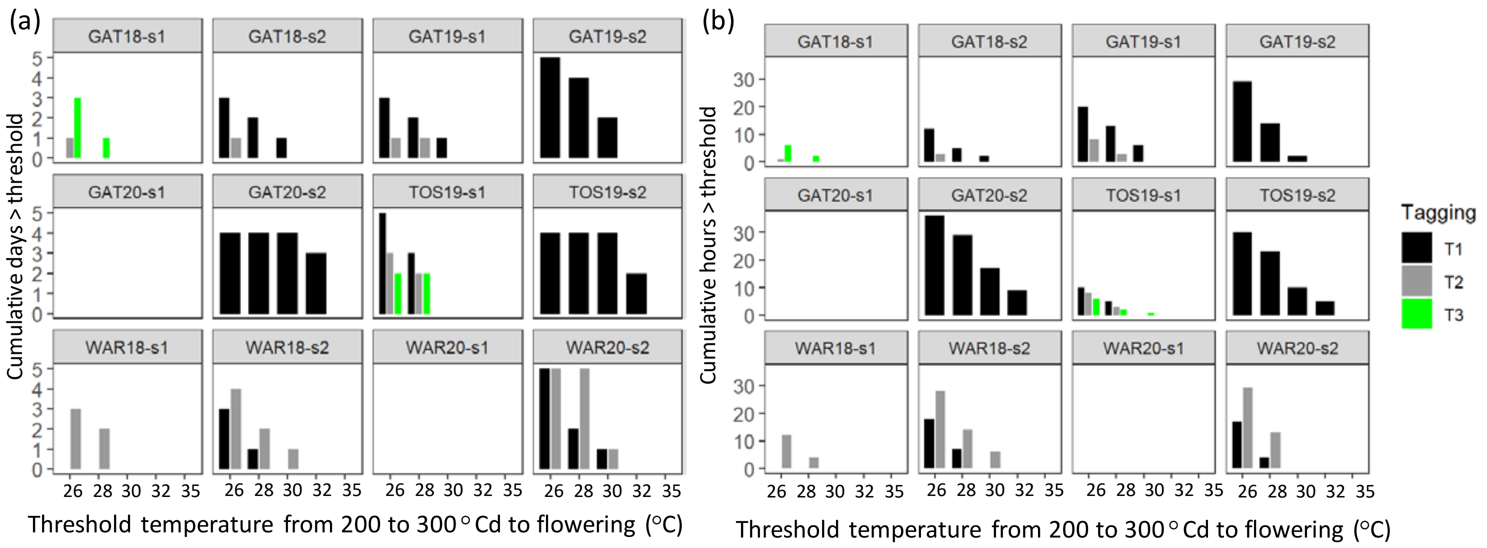


**Figure S2: Cumulative pre-flowering (200-300°Cd before flowering) heat days (a) and hours (b) with temperatures above threshold temperature experienced by wheat crops in the studied environments.** T1, 1st cohort of stems tagged at flowering, T2: 2nd cohort of stems tagged at flowering, T3: 3rd cohort of stems tagged at flowering.

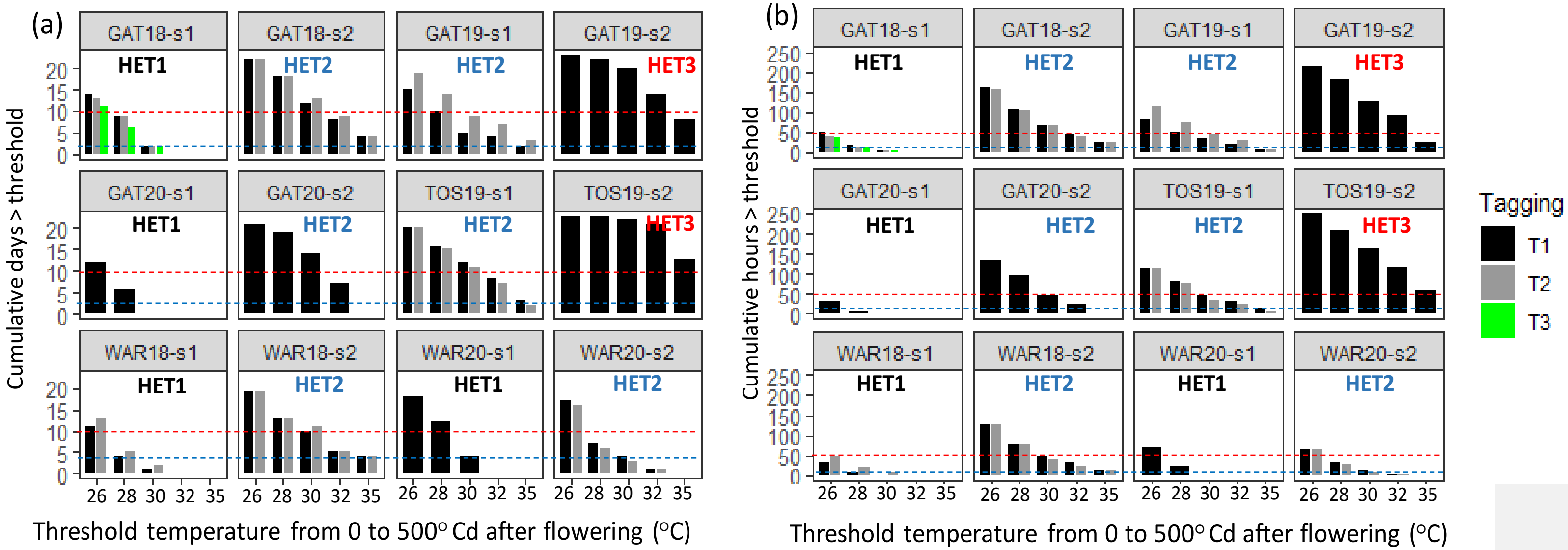


**Figure S3: Cumulative post-flowering (0-500°Cd after flowering) heat days (a) and hours (b) with temperatures above threshold temperature experienced by wheat crops in the studied environments.** T1, 1st cohort of stems tagged at flowering, T2: 2nd cohort of stems tagged at flowering, T3: 3rd cohort of stems tagged at flowering.

Based on post-flowering heat environments were classified into three heat environment types (HETs, Fig. S3); heat environment type 1 (HET1, black label) corresponds to environments with 0 days or hours of maximum $T_{thresh}$ > 32°C between 0 and 500°Cd after flowering, while heat environment type 2 (HET2, blue coloured label) corresponds to 1-9 days or 1-49 of $T_{thresh}$ > 32°C during the same period. Heat environment type 3 (HET3, red label) had more than 10 days or 50 hours of $T_{thresh}$ > 32°C between 0 and 500°Cd after flowering.
